# Historical Demography and Climate Driven Range Shifts in the Blue-spotted Salamander Under the Climate Change Scenarios

**DOI:** 10.1101/2022.09.26.509446

**Authors:** Utku Perktaş, Can Elverici, Özge Yaylali

## Abstract

This study integrates phylogeography with distributional analysis to understand the demographic history and range dynamics of a limited dispersal capacity amphibian species, Blue-spotted Salamander (*Ambystoma laterale*), under several climate change scenarios. For this we used an ecological niche modeling approach, together with Bayesian based demographic analysis, to develop inferences regarding this species’ demographic history and range dynamics. The current model output was mostly congruent with the present distribution of the Blue-spotted Salamander. However, under both the Last Interglacial and the Last Glacial Maximum bioclimatic conditions, the model predicted a substantially narrower distribution than the present. These predictions showed almost no suitable area in the current distribution range of the species during almost the last 22.000 y before present (ybp). The predictions indicated that the distribution of this species shifted from eastern coast of northern North America to the southern part of the current distribution range of the species. The Bayesian Skyline Plot analysis, which provided good resolution of the effective population size changes over the Blue-spotted Salamander history, was mostly congruent with ecological niche modeling predictions for this species. This study provides the first investigation of the Blue-spotted Salamander’s late-Quaternary history based on ecological niche modeling and Bayesian-based demographic analysis. In terms of the main result of this study, we found that the species’ present genetic structure has been substantially affected by past climate changes, and this species has reached current distribution range almost from nothing since the Last Glacial Maximum.

## Introduction

The effects of late-Quaternary climate changes on North American biodiversity are considered to be dramatic (Pielou, 1991). Recent distribution patterns must have occurred from glacier-free regions during the Last Glacial Maximum (LGM). These recolonization events generally took place rapidly to preglaciated regions (G. Hewitt, 2000). It is generally accepted that these refugia were located in the south because evidence from palynological studies (Webb *et al*., 2003) shows that large changes in plant communities were also occurring rapidly. Some recent studies have also shown that the distribution areas of some North American species have changed almost completely, and their distribution has shifted from south to north due to the changing climate (Perktaş and Elverici, 2020). Therefore, many northern taxa have lower genetic diversity than their southern counterparts. This is probably due to the fast post-glacial re-colonization, partial extinction, and the fast colonization events in a short time (Hewitt, 1996). Until recently, this low genetic variability has hampered our ability to detect the unique genetic structure that can be found in taxa at higher latitudes.

The evaluation of mitochondrial DNA (mtDNA) diversity gives an opportunity to understand species’ demographic history from the recent past to the present (Freeland, 2005) Integrating mtDNA diversity with distributional analyses offers novel opportunities to understand such complex biogeographic stories (e.g., Klicka *et al*., 2011). The Blue-spotted Salamander (*Ambystoma laterale*) has one of the northern-most distributions in North America with very limited dispersal ability (Conant and Collins, 1991). During the Last Glacial Maximum (LGM, approx. 22,000 years before present), ice coverage extended over almost the entire current distribution range of the Blue-spotted Salamander (Fig. 1). Hence, in this paper, we aim to develop an integrative biogeographic survey on the Blue-spotted Salamander to evaluate its demographic history under the climate change scenarios. This study can be considered as an integration and continuation of the work of Demastes *et al*. (genotype; 2007) with distributional projections derived from ecological niche models (ENMs).

**Figure 1.**
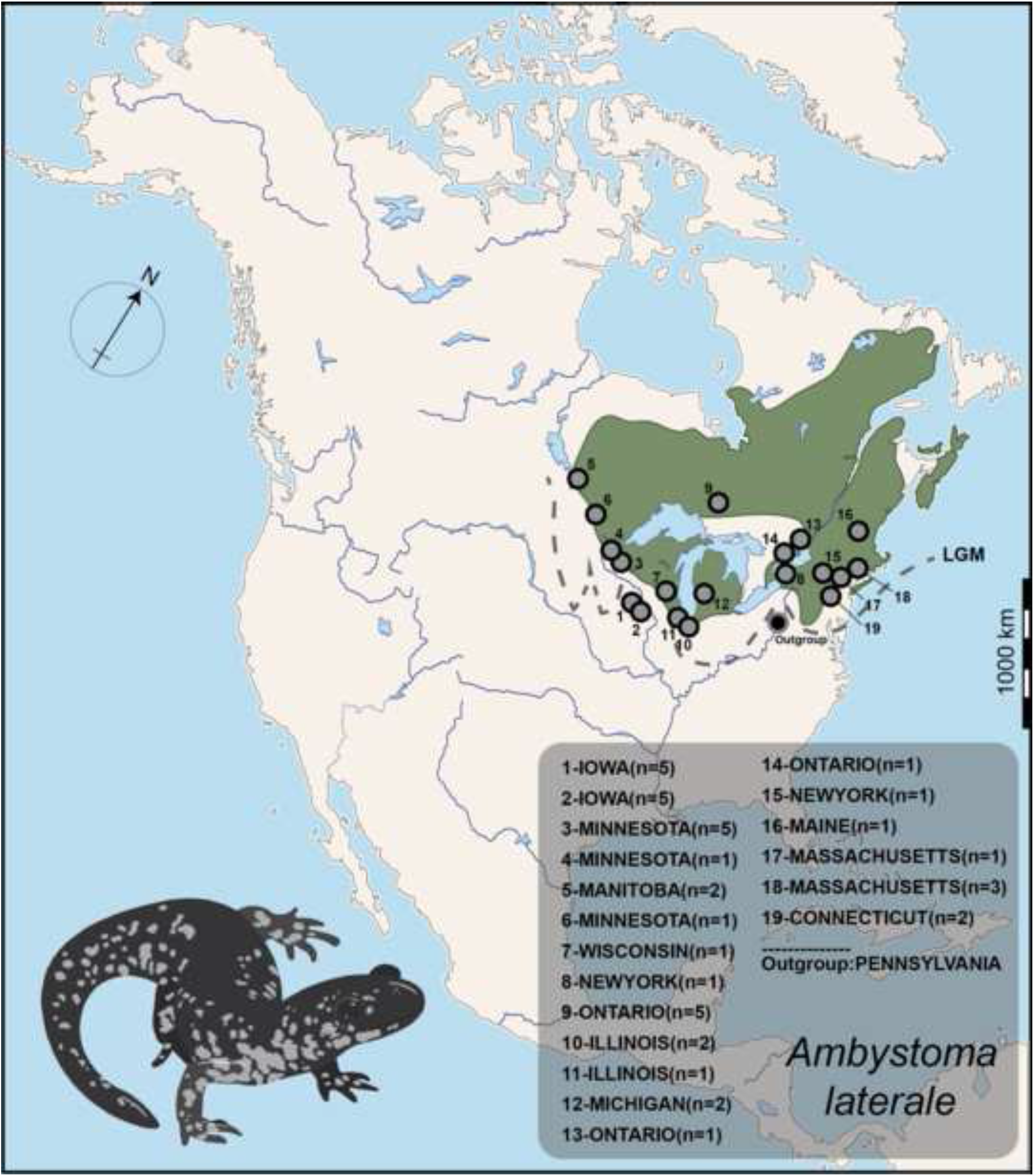
Approximate distribution range of the Blue-spotted Salamander. Sampling localities for ingroup taxa (1–19) are indicated based on Demastes *et al*. (2007). Dashed line depicts the approximate extent of the last glacial maximum

## Methods

### ECOLOGICAL NICHE MODELLING

Input data – We analyzed species occurrence data from GBIF (www.gbif.org), ranging from 1964 to 2020 (n = 2561 after 781 duplicated occurrence records removed), before checking for sampling bias and spatial autocorrelation (Brown, 2014) for occurrence records. We spatially filtered all records to eliminate multiple records, leaving single 20 km records across the species’ distribution. For this we considered the low dispersal capacity of the Blue-spotted Salamander (see Ryan and Calhoun, 2014; Vanek *et al*., 2019 for details). This yielded 553 unique occurrence records for ecological niche modelling.

We downloaded bioclimatic data from the WorldClim database (Hijmans *et al*., 2005, http://www.worldclim.org) for the Last Interglacial, three global climate models (CCSM4, MIROC-ESM, and MPI-ESM-P) for the Last Glacial Maximum (∼22 kybp), the mid-Holocene (∼6 kybp), the present (∼1960-1990), and future conditions based on the RCP4.5 and RCP8.5 greenhouse gas scenarios (2050 and 2070) at a spatial resolution of 2.5 arc-minutes. Bioclimatic data included 19 bioclimatic variables derived from monthly temperature and precipitation values. Previous studies (Campbell *et al*., 2015; Escobar *et al*., 2014) detected a number of apparent artifacts in some of the climate datasets: mean temperature of the wettest quarter, mean temperature of driest quarter, precipitation of the warmest quarter, and precipitation of the coldest quarter (BIO8–9, BIO18–19, respectively). Therefore, we excluded these variables from the pool of selected variables to build a model (for details, also *see* Simoes *et al*., 2020). Since the Blue-spotted Salamander is one of the rarest amphibian species in northeastern North America (Fig. 1; Ryan and Calhoun, 2014), all variables were masked to include all North America (-170° to 13° W and - 50° to 84° N). We then inspected correlations between these bioclimatic variables to produce three different climatic data sets based on different inter-variable correlation coefficients (0.6, 0.7, 0.8, and 0.9). These included mean diurnal range (BIO2), isothermality (BIO3), maximum temperature of warmest month (BIO5), annual temperature range (BIO7), precipitation of wettest month (BIO13) and driest month (BIO14) and precipitation seasonality (BIO15).

### DEMOGRAPHIC HISTORY

We applied the Bayesian Skyline Plot (BSP) method, implemented in BEAST version 1.10.4 (Suchard *et al*., 2018), to explore the Blue-spotted salamander demographic history. Since earlier studies have not reported any structure (Demastes *et al*., 2007), we combined all mtDNA sequences before running the BSP analysis, and this approach made the demographic history more comparable with ecological niche modelling. Before the BSP runs, the best-fit substitution models were identified for the mtDNA control region sequences in MEGA X (Kumar *et al*., 2018). These were the Hasegawa-Kishino-Yano (HKY, AICc = 1775.424) for the control region. Multiple independent Bayesian Skyline Plot runs were performed using the following parameters: linear models, 10 million steps, parameters sampled every 1000 steps, and a burn in of 10%. For the control region sequences, we used the strict clock model with a default mutation rate under normal prior distribution [for vertebrates, the widely-used 2%-6% substitutions/site/million years (Allio *et al*., 2017; for different examples, *see* also Brito, 2005; Pereira and Baker, 2006)]. The effective sample size values of the parameters were over 200 for each run.

## Results

We evaluated 2108 candidate models using combinations of 31 feature classes, 17 regularization multipliers, and 4 climatic data sets. The best model for the Blue-spotted Salamander was provided by the third climatic data set which was had a correlation threshold of 0.8 (Set 3: BIO3, BIO5, BIO7, BIO13, BIO14 and BIO15), which was significantly different from random (P < 0.001) and had the lowest Akeike information criteria set. The model had a regularization multiplier of 10 and included one feature class (threshold). Projections for past, present, and future performed better than a random prediction with moderate training AUC = 0.708 with small standard deviation (sd = 0.022) that indicated the robust model performance. Three bioclimatic variables contributed the most to the model (together 66.5%): BIO5 (30.5%), BIO15 (19%) and BIO7 (17%).

Under present bioclimatic conditions, the model’s predictions were mostly congruent with the present and recent historical distribution of Blue-spotted Salamander (*see* Fig.1 for the present distribution, also see IUCN, 2015). The model primarily predicted areas of high suitability across habitats for the species in North America. Under the Last Glacial Maximum bioclimatic conditions, the model predicted substantially narrower distribution than the present and mid-Holocene (Fig. 2). Interestingly, predictions for the Last Glacial Maximum indicated almost no distribution in the east-coast of North America. However, predictions for the Last Interglacial indicated a distribution east coast of North America. For both 2050 and 2070, the model predicted that the range will most likely shift slightly northward with a wider distribution than either the past or present (Fig. 2).

**Figure 2.**
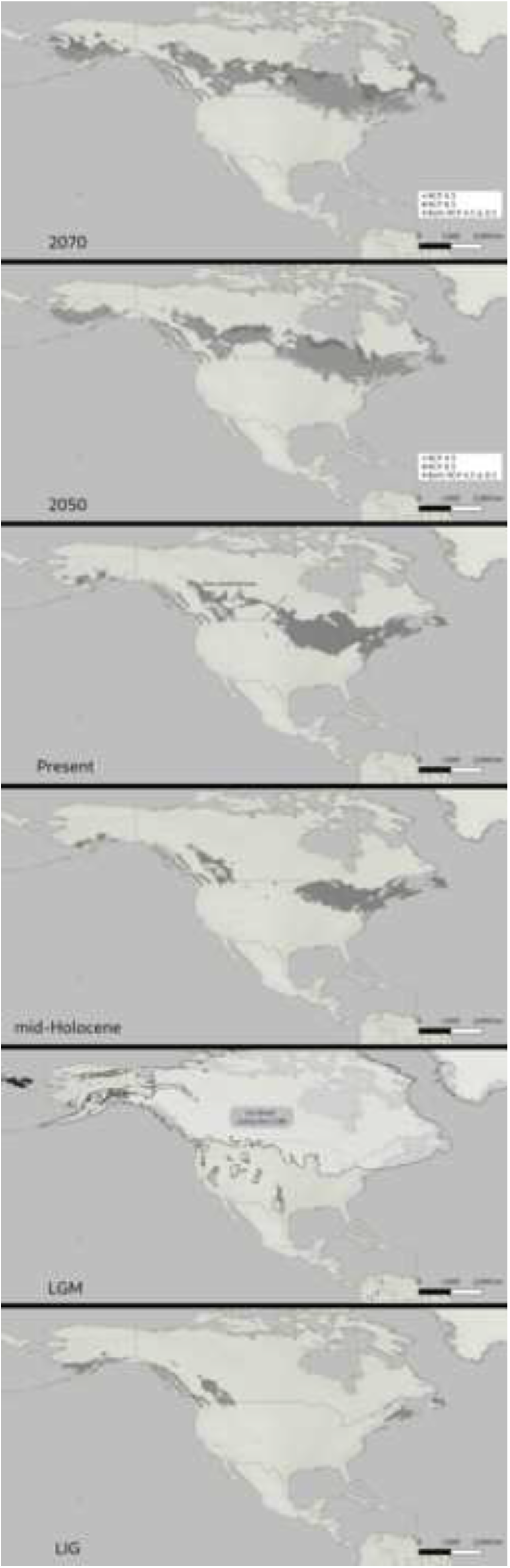
Ecological niche model-based distributional predictions for the Blue-spotted Salamander under the different bioclimatic conditions [*i*.*e*. the LGM and the Future (2050 and 2070)]

Based on a strict molecular clock (mean 2%-4% substitutions/site/million years), the BSP result provided a good resolution of the effective population size changes over the Blue-spotted Salamander history (Fig. 3). The BSP indicated a recent demographic expansion starting after the Last Interglacial based on both mutation rates (starting approximately after 60,000 y before present).

**Figure 3.**
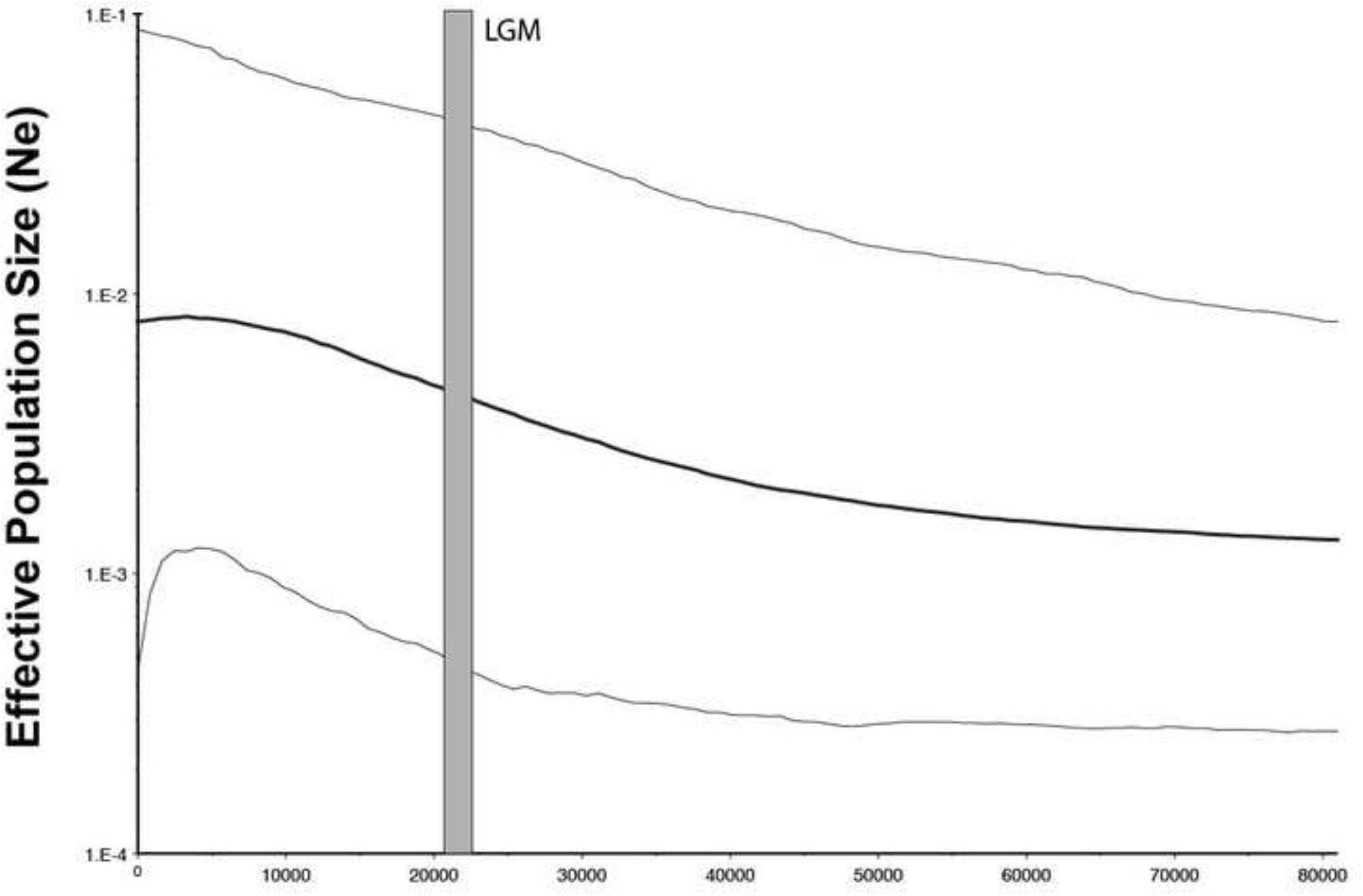
The effective population size fluctuation of the Blue-spotted Salamander based on the Bayesian Skyline Plot analysis

## Discussion

This study focuses on the dynamics of range shifts of the Blue-spotted Salamander from the past, the present and the near future under climate change scenarios. Accordingly, this study is therefore a continuation of the work of Demastes *et al*. (2007) with the first investigation of the late-Quaternary history of the Blue-spotted Salamander based on ecological niche modelling and Bayesian based demographic analysis.

The Blue-spotted Salamander showed substantially low-level genetic diversity based on 534 nucleotides of non-coding mtDNA gene region (Demastes *et al*., 2007). During the Last Glacial Maximum, north-eastern North America was almost completely covered by ice (Pielou, 1991) and there is no fossil evidence to suggest that salamanders were present south of the ice sheet. Therefore, lack of mtDNA differentiation in the Blue-spotted Salamander across its distribution range suggests that populations arose recently (after the Last Glacial Maximum) from a relatively homogeneous ancestral population. Phylogenetic relationships among haplotypes (*see* Fig. 2 in Demastes *et al*., 2007) indicated most haplotypes are closely related, yet are geographically localized. The western clade separated from the east coast and central clades with high bootstrap values. All these signals suggest that populations are historically connected, but most probably due to behavioral reasons, populations are sundered by firm, and it supports recent and continuing genetic differentiation. This situation can be discussed as the absence of a long-term biogeographical barrier, but depends on the limitations in dispersal capacity of this species. The current pattern is compatible with phylogeographic category III (shallow gene tree, allopatric lineage) specified by Avise *et al*. (1987) and (Avise, 2000).

Empirical examples that support this phylogeographic pattern from different species that spread out of the glacial line, along the southeastern coastline of North America (eg Deer Mouse, *Peromyscus polionotus*, Avise *et al*., 1979, 1983), South America (White-Tailed Deer, *Odocoileus virginianus*, Moscarella *et al*., 2003) has been published. However, it is well known that northeast North America has undergone remarkable topographic change because of past climatic changes, including climatic fluctuations during the last 130.000 ybp, and repeated expansion /recession of continental ice sheets during especially 22.000 ybp, and therefore habitat changes in the same time period. All these events have had dramatic effects on the genetic structuring of flora and fauna in the region (*e*.*g*. Pielou, 1991); and hence the genetic structure of Blue-spotted Salamander showed tight association with past climate change events.

The past distributional predictions of the Blue-spotted Salamander indicated substantial range-shifts from the Last Interglacial to the Present. The predicted Last Interglacial range limited to the east coast of northern North America, then the predicted Last Glacial Maximum range limited to the southern range of the distribution range of the species. Therefore, the range of Blue-spotted Salamander slided from east to south, and expanded to the north and reached the current distribution. Species looks like it experienced an almost complete extinction in its present distribution range during the Last Glacial Maximum. According to Lindsay *et al*. (2016), habitat suitability for the Blue-spotted Salamander in the Last Glacial Maximum, as ecological niche modelling approach predicts, is mostly Taiga and partly Montane Mosaic. IUCN states that there are two main suitable habitats for the species, forests and wetlands. Suitable forests are boreal (taiga) and temperate forests, so the Last Glacial Maximum vegetation model agrees with our niche modelling results. The Southern Appalachian Region was found to be a refugium for many species (Hewitt, 2004). However, this type of biogeographic pattern has not been reported for any terrestrial vertebrate species so far although some similar examples have been reported for widespread North American vertebrates (*e*.*g*. Klicka *et al*., 2011; van Els *et al*., 2012; Barrowclough *at al*., 2018; Perktaş and Elverici, 2020). The Bayesian Skyline Plot analysis showed a population increase before the Last Glacial Maximum in the ice free areas in North America, and this makes this study outputs interesting. The species, however, reached its present distribution gradually during the Holocene. Therefore, the species’ demographic history supports the ecological niche modelling results. In contrast to phylogeographic research on other vertebrate species (*e*.*g*. Sharp-tailed Grouse), we found no evidence of a large refugium in the ice free range of North America for this species. However, the Blue-spotted Salamander has almost completely changed its distribution since the last glacial period; that is, this species has reached current distribution range almost from nothing since the Last Glacial Maximum.

## Acknowledgment

We thank the remaining members of the Biogeography Research Laboratory for their support and assistance in various phases of this project.

## Author Contributions

UP conceived the study; UP developed methods and UP, CE, ÖY analyzed data; CE visualized the distributional predictions, UP wrote the paper with discussion with CE and ÖY. All authors read and approved the final manuscript.

